# Transcription closed and open complex formation coordinate expression of genes with a shared promoter region

**DOI:** 10.1101/842484

**Authors:** Antti Häkkinen, Samuel M. D. Oliveira, Ramakanth Neeli-Venkata, Andre S. Ribeiro

## Abstract

Many genes are spaced closely, allowing coordination without explicit control through shared regulatory elements and molecular interactions. We study the dynamics of a stochastic model of a gene-pair in a head-to-head configuration, sharing promoter elements, which accounts for the rate-limiting steps in transcription initiation. We find that only in specific regions of the parameter space of the rate-limiting steps is orderly co-expression exhibited, suggesting that successful cooperation between closely spaced genes requires the co-evolution of compatible rate-limiting step configuration. The model predictions are validated by *in vivo* single-cell, single-RNA measurements of the dynamics of pairs of genes sharing promoter elements. Our results suggest that, in *E. coli*, the kinetics of the rate-limiting steps in active transcription can play a central role in the dynamics of pairs of genes sharing promoter elements.

## I. INTRODUCTION

Closely-spaced gene-pairs abound in genomes of all life forms, from human [1, 2] to prokaryotes [3, 4]. Further, they are highly conserved [2, 5], suggesting that they yield functionalities with selective advantages.

Gene-pairs can be arranged head-to-head (transcriptionally divergent), with their transcription start sites (TSS) closely located, sharing promoter elements such as transcription factor binding sites [1, 2, 4]. Head-to-tail (tandem) and tail-to-tail (convergent) overlapping genepairs are also found, allowing interference between RNA polymerases (RNAP) [6] and/or with transcription factors [7, 8]. Each configuration can vary in several parameters, such as distance between TSSs, which affect transcription of the component genes [4, 9–11], allowing co-regulation without explicit control mechanisms. The multitude of naturally occurring configurations suggests that each yields distinct selective advantages.

While some configurations have been identified and their ubiquity established by models and measurements [2, 11–13], the range of possible behaviors and advantages as a gene regulation mechanism remain largely uncharacterized. Such characterization would benefit understanding the array of tasks that organisms such as *Escherichia coli* perform using closely-spaced promoters, as opposed to individual genes or genes connected by transcription factors.

One aspect not yet considered is the existence of multiple rate-limiting steps during transcription initiation [14, 15]. As only some of these steps are physically involved in the gene-pair interactions, we expect the nature of the rate-limiting steps of each promoter to affect the dynamics of closely-spaced configurations. Importantly, the durations of the open complex formation of a strong and a weak promoter can differ from little to up to two orders of magnitude [16] and live cell single-RNA measurements suggest that different promoters are rate-limited at different stages of transcription initiation [15, 17–20]. As such, it is plausible that promoters whose initiation kinetics are similar in mean duration but whose rate-limiting step structures differ will feature different dynamics in the bidirectional configuration.

Here, we study the dynamics of a stochastic model of a gene-pair in a head-to-head configuration sharing promoter elements (the most common closely-spaced gene-pair configuration [2, 5]) as a function of the rate-limiting step configuration of each gene. We analyze the models using analytical stochastic methods. Next, we validate the main findings by performing time-lapse microscopy measurements of individual genes and in pairs of genes sharing promoter elements, at the single-RNA level, in live *E. coli*.

## II. METHODS

### A. Models

Transcription in *E. coli* starts when an RNAP, recruiting the appropriate s-factor, specifically binds to a promoter. This creates a closed complex of the RNAP and DNA, which can require several trials before stabilizing [21]. In strong promoters, this step is nearly irreversible [22]. The virtually irreversible open complex formation follows, consisting of e.g. DNA unwinding and compaction [23] and the RNAP clamp assembly [24].

We assume a variant of a model of transcription initiation of the overlapping promoters of the galactose operon in the absence of cAMP-CRP [3]. The transcribed promoter stochastically is selected based on the relative affinities between the two promoters and the RNAP, encoded in the forward rates of the closed complex formation of each promoter. After the selection, the remaining steps of transcription initiation occur at the promoter region [14]. The following stochastic chemical [25, 26] reactions are used to model this:

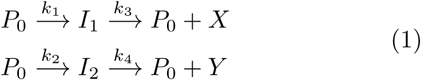

where *P*_0_ represents a free promoter (unoccupied by an RNAP), *I*_1_ and *I*_2_ represent intermediate transcriptional complexes committed to transcribing genes 1 and 2, respectively, and *X* and *Y* represent the messenger RNA products (or, if they closely follow [27], proteins) of genes 1 and 2, respectively. A schematic is provided in Fig 2A, and a thorough analysis in the supplement.

**FIG. 1.**
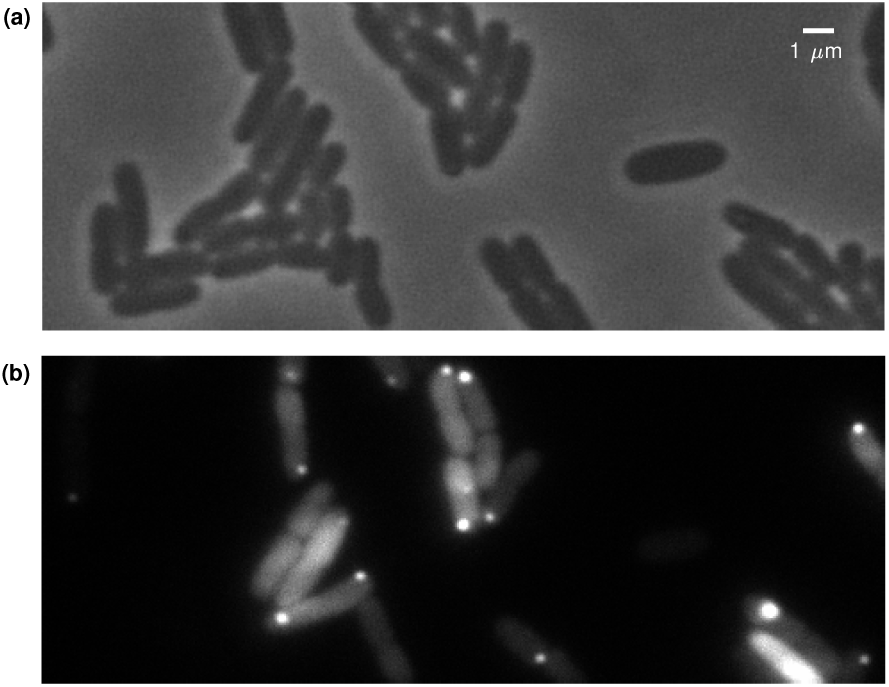
Example images of live *E. coli* expressing GFP-tagged RNAs. (A) Phase contrast image of the live *E. coli* with the P_lacO3O1-tetA_ construct taken after 1 hour of induction with 1 mM IPTG induction at 37 °C. (B) HILO image visualizing the abundant GFP inside the same *E. coli* cells and the target RNA bound by an array of GFPs appearing as bright spots.

**FIG. 2.**
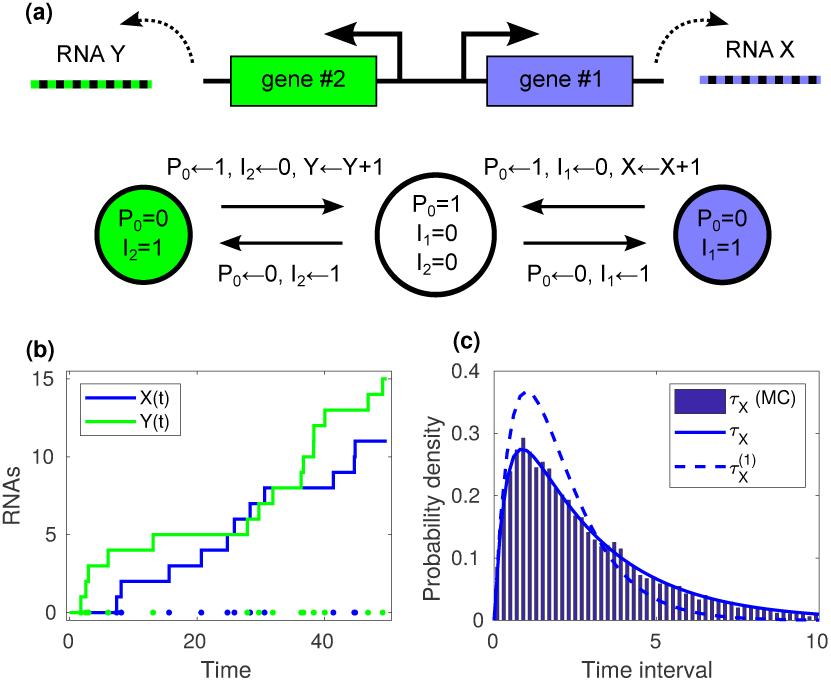
Model schematic and simulated examples. (A) Schematic of gene-pair in a head-to-head configuration: genes 1 and 2 produce RNAs X and Y, respectively. The shared promoter can be in three-states: free, or occupied for transcription initiation of gene 1 or 2. (B) Produced RNA numbers over time in a single Monte Carlo simulation. The dots denote the moments when RNAs were produced. (C) Distribution of intervals between consecutive productions of *X* in 10,000 simulations. The parameter values are (*k*_1_, *k*_2_, *k*_3_, *k*_4_)=(1, 1, 1, 1). Here, *τ*_*X*_ and *τ*_*X*_^(1)^ have a mean (variance) of 3 (7) and 2 (2), respectively.

If the genes did not share promoter elements, the intervals between productions of *X* (gene 1) would be [20]:

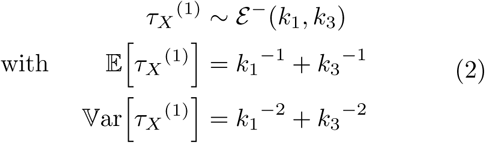

where *ε*^−^(*λ*_1_,…, *λ*_*n*_) represents a hypoexponential distribution with rates *λ*_1_, …, *λ*_*n*_. Similarly, the production intervals for gene 2 would be *τ*_*Y*_^(1)^ ∼ *ε*^−^(*k*_2_, *k*_4_).

These distributions are of low noise, as measured by the coefficient of variation (standard deviation over the mean), as this quantity equals unity for Poissonian production (exponential production intervals). The noise is determined by the ratio of *k*_1_ and *k*_3_. Regardless of the mean, it is minimized for steps of equal duration and maximized when a single step is rate-limiting. The dynamics of an individual gene is unaffected by the step order (i.e. interchanging *k*_1_ and *k*_3_ has no effect on *τ*_*X*_^(1)^).

Regardless of the configuration, the mean and variance of the production intervals are linked to that of the produced RNAs. In the long-term (infinite time), the mean and variance of produced RNA per unit time are [28]:

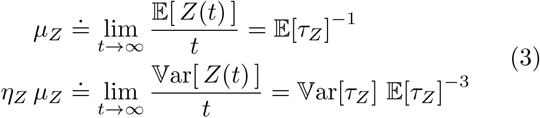

i.e. the mean number of RNAs produced per unit time (*μ*_*Z*_) equals the inverse interval mean, while the Fano factor (variance over the mean) of the RNA numbers (*η*_*Z*_) equals the squared coefficient of variation of the production intervals. The cell phenotype is also affected by other processes, such as RNA degradation and dilution due to cell division. Regardless, the mean and noise of the produced RNA numbers are directly linked to the phenotype (details in supplement) [29], so we expect our results to hold qualitatively in the presence of other processes.

### B. Cells, plasmids, chemicals, and growth conditions

We used *E. coli* strain BW25113 (*lacI* ^+^ *rrnB*_T14_ Δ*lacZ* _WJ16_ *hsdR514* Δ*araBAD*_AH33_ Δ*rhaBAD*_LD78_) [30], which contains the constitutive promoters P_lacI+_ and P_araC_ producing, respectively, LacI repressors [31] and AraC repressors. As this strain does not contain the *tetR* gene responsible for encoding TetR repressors, any gene downstream to a P_tetA_ promoter is expressed constitutively.

We constructed five target systems on a single-copy pBELO plasmid. The first plasmid features the P_lacO3O1_ promoter controlling the production of an RNA molecule coding for a red fluorescent mCherry protein followed by 48 binding sites for the MS2-GFP protein (mCherry-48BS). The other four systems are modified versions of the first, with the P_lacO3O1_ promoter being replaced by the following promoters: (i) P_BAD_ promoter; (ii) P_lacO3O1-tetA_ dual-tandem promoter; (iii) P_lacO3O1-BAD_ dual-tandem promoter; and (iv) P_lacO3O1-lacO3O1_ dual-bidirectional promoter. All strains aside from its target system also contain either a medium-copy plasmid pZA25 with the reporter gene P_ara_-MS2-GFP or a low-copy plasmid pZS12 with the reporter gene P_lac_-MS2-GFP. These plasmids are responsible for producing the fusion protein MS2-GFP, both producing an abundance of MS2-GFP when activated as detailed below. The reporter plasmids were generously provided by Orna Amster-Choder (Hebrew University of Jerusalem, Israel) [32], and Philippe Cluzel (Harvard University, USA) [33], respectively. The activity of the promoters P_lacO3O1_, P_lacO3O1-tetA_, and P_lacO3O1-lacO3O1_ is regulated by the repressor LacI and the inducer isopropyl β-D-1-thiogalactopyranoside (IPTG). Meanwhile, the activity of P_BAD_ is regulated by the repressor AraC and the inducer L-arabinose. Finally, the activity of P_lacO3O1-BAD_ is regulated by both repressors (LacI and AraC) and both inducers (IPTG and L-arabinose).

Cells were grown overnight in lysogeny broth (LB) medium supplemented with appropriate antibiotics (34 μg/ml of chloramphenicol, 50 μg/ml of ampicillin, and 50 μg/ml of kanamycin) with shaking at 250 rpm. We made subcultures, by diluting the stationary-phase culture into fresh M9 medium supplemented with glycerol (0.4% final concentration) and the appropriate antibiotics. Cells were left in the incubator until reaching OD_600_ of about 0.25. For the pZA25-P_ara_-MS2-GFP reporter plasmid activation, 0.4% of L-arabinose was added to the culture, which was then incubated at 37 °C for 60 minutes. Cells containing the pZS12-P_lac_-MS2-GFP reporter plasmid were incubated in the same way and were activated with 1 mM IPTG. Next, for the activation of P_lacO3O1_, P_lacO3O1-tetA_, and P_lacO3O1-lacO3O1_ target plasmids, specific concentrations of IPTG (either 5 μM or 1 mM) were added to the culture. For activating the P_BAD_ or P_lacO3O1-BAD_ target plasmids, 0.1% of L-arabinose was added. For the latter, similar concentrations of IPTG (5 μM or 1 mM) were added as well. Inducer-activated cells were then left in the incubator for 90 minutes, prior to microscopy observation.

### C. Microscopy and Image analysis

Cells were visualized using a Nikon Eclipse (Ti-E, Nikon) inverted microscope equipped with a 100× Apo TIRF (1.49 NA, oil) objective. Cells and fluorescent spots within were imaged by Highly Inclined and Laminated Optical sheet (HILO) microscopy, using an EMCCD camera (iXon3 897, Andor Technology), a 488 nm argon laser (Melles-Griot), and an emission filter (HQ514/30, Nikon). Phase-contrast images were acquired by a CCD camera (DS-Fi2, Nikon). The software for image acquisition was NIS-Elements (Nikon, Japan). An example of each channel is shown in Fig 1.

We performed time-lapse fluorescence and phase-contrast imaging of the cells (the latter for cell segmentation and lineage construction). For this, 8 μl of cells were placed on a microscope slide between a coverslip and a M9 glycerol agarose gel pad. During image acquisition, cells were constantly supplied with fresh media containing IPTG and L-arabinose, at the same concentration as when in liquid culture, by a micro-perfusion peristaltic pump (Bioptechs) at 0.3 ml/minute. Images were captured for 5 hours, once per minute in the case of fluorescence and once per 5 minutes in the case of phase-contrast. During image acquisition, cells were kept in a temperature-controlled chamber (FCS2, Bioptechs) at optimal temperature (37 °C).

Time series microscopy images were processed as in [34] by, first, aligning consecutive images so as to maximize the cross-correlation of fluorescence intensities. Next, we annotated manually the region occupied by each cell in the time series. Afterwards, the location, dimension, and orientation of each cell in each frame is obtained by principal component analysis, assuming that fluorescence inside the cell is uniform [35]. Cell lineages were then extracted using CellAging, based on overlapping areas in consecutive frames [35]. Next, the intensity of each cell is fit with a surface (quadratic polynomial of the distance from the cell border) in least-deviations sense [36]. This surface represents the cellular background intensity which is subtracted to obtain the foreground intensity. Next, the foreground intensity is fit with a set of Gaussian surfaces, in least-deviations sense, with decreasing heights until the heights are in the 99% confidence interval of the background noise (estimated assuming a normal distribution and using median absolute deviation) [36]. The Gaussians represent fluorescent RNA spots, and the volume under each represent the total spot intensity. Finally, as MS2-GFP-tagged RNA lifetimes are much longer than cell division times [37], the cellular foreground intensity will be an increasing curve, with each jump corresponding to the appearance of a novel tagged RNA. The moments when a jump occurs are estimated using a specialized curve fitting algorithm [18]. The intervals between jumps in individual cells correspond to time intervals between consecutive RNA production events.

## III. RESULTS AND DISCUSSION

### A. Analytical distributions of production time intervals

From the perspective of the production kinetics of *X* alone, the reaction system of Eq (1) is equivalent to:

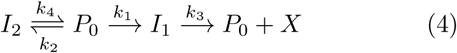

which is potentially a highly noisy process [20, 38]. While the expression of gene 1 might not be noisy on its own, its expression is perturbed by the transcription machinery occupying the shared promoter region for expression of gene 2, introducing (random) temporal gaps in the expression.

Let 𝒢(·) denote the distribution of consecutive productions of *X* in Eq (4). The mean and variance of the time intervals between the productions of *X* are given by [20]:

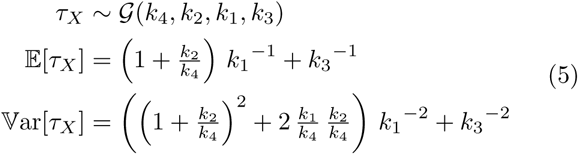

while, due to the symmetry of the model, the production intervals of *Y* are *τ*_*Y*_ ∼ 𝒢(*k*_3_, *k*_1_, *k*_2_, *k*_4_).

By comparing Eq (2) with Eq (5) we find that, regardless of the parameters, in a bidirectional configuration, the mean and variance of the time intervals between RNA productions of each gene are increased. Consequently, while *τ*_*X*_^(1)^ is always sub-Poissonian, *τ*_*X*_ can exhibit either sub- or super-Poissonian behavior.

RNA production according to the model is exemplified in Fig 2B, and the expected interval distribution in Fig 2C. While the production intervals of each gene are often somewhat regular, as indicated by the bulk of the distribution, large outliers are present due to the temporal gaps, which coincide with the transcriptional activity of the other gene (cf. Fig 2B).

As the marginals shown in Eq (5) fail to capture the co-expression of the two genes, further analysis is necessary. The time between consecutive productions by either gene, i.e. a jump in *X*(*t*) + *Y* (*t*), is (detailed in the supplement):

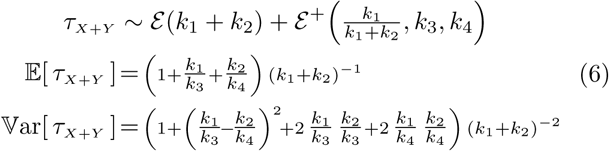

where *ε*(*λ*) is an exponential distribution with rate *λ* and *ε*^+^(*p*_1_, …, *p*_*n*−1_, *λ*_1_, …, *λ*_*n*_) is a hyperexponential distribution with mixing probabilities *p*_1_, …, *p*_*n*_ and rates *λ*_1_, …, *λ*_*n*_. Again, this distribution can feature either sub- or super-poissonian behavior, depending on its parameter values. By combining Eq (3), (5), and (6), one can determine the asymptotic covariance and the (Pearson) correlation *ρ*_*XY*_ between the produced RNA numbers *X*(*t*) and *Y* (*t*) (detailed in the supplement).

### B. Noise and correlation in the transcription kinetics of genes in a head-to-head configuration

Based on the above, we first analyzed how the noise and correlation in the transcription kinetics of a head-to-head configuration depends on the dynamics of the individual genes. For this, the parameterization *λ, q*_12_, *q*_13_, *q*_24_ was found to be insightful. Here, *λ* ≐ *μ*_*X*+*Y*_ is a timescale parameter (mean total production rate) and *q*_*ij*_ ≐ *k*_*i*_ */ k*_*j*_ denote ratios of rates of two reactions. Further, *q*_12_ controls the bias, i.e. the expression ratio of each gene: for large (small) *q*_12_, gene 1 (gene 2) is expressed more frequently. Finally, *q*_13_ and *q*_24_ control the relative durations of closed and open complex formation, which equal 1 */* (1+*q*_13_) and *q*_13_ */* (1+*q*_13_), respectively, for gene 1. Specifically, if *q*_13_ > 1 (*q*_13_ < 1), then *k*_1_ > *k*_3_ and the gene is limited at the open (closed) complex formation.

The RNA number means are controlled by the bias and the scale: *μ*_*X*_ = *λ*^−1^ *q*_12_ */* (1+*q*_12_) and *μ*_*Y*_ = *λ*^−1^ */* (1+*q*_12_). As such, the stage at which the transcription kinetics of each gene is rate-limited does not affect the mean number of produced RNAs. Meanwhile, the noise and correlation exhibit complex behavior, which can be divided into a few regions. The regions and their properties are shown in Table I (and Table S-I). The noise of each gene and the correlation coefficient are shown in Fig 3 and Fig 4A, and their analytical forms in the supplement.

**TABLE I.** Noise and correlation in RNA production kinetics in the different regions of the parameter space of head-to-head configuration. Here, ∼1^−^ (∼1^+^) denotes weakly sub-(super-) Poissonian behavior (noise of about 1), while 1 denotes that both behaviors are possible. Finally, < 1^*^ indicates that < 1 holds at least for one of the genes, possibly for both.

| Region | Condition | Noise $\eta_X$ | Noise $\eta_Y$ | Corr. $\rho_{XY}$ |
| --- | --- | --- | --- | --- |
| A | $q_{24} > 1, q_{24} > q_{13}$ | $> 1$ | $\sim 1^-$ | $> 0$ |
| A | $q_{13} > q_{24}, q_{13} > 1$ | $\sim 1^-$ | $> 1$ | $> 0$ |
| B | $q_{13}, q_{24} < 1$ | $\sim 1$ | $\sim 1$ | $\sim 0^-$ |
| C | $q_{13} \sim q_{24} > 1$ | $< 1^*$ | $< 1^*$ | $< 0$ |
| D | $q_{13} \sim 1, q_{24} < 1$ | $< 1$ | $\sim 1^+$ | $< 0$ |
| D | $q_{13} < 1, q_{24} \sim 1$ | $\sim 1^+$ | $< 1$ | $< 0$ |
| E | $q_{13} \sim q_{24} \sim 1$ | $< 1^*$ | $< 1^*$ | $< 0$ |

**FIG. 3.**
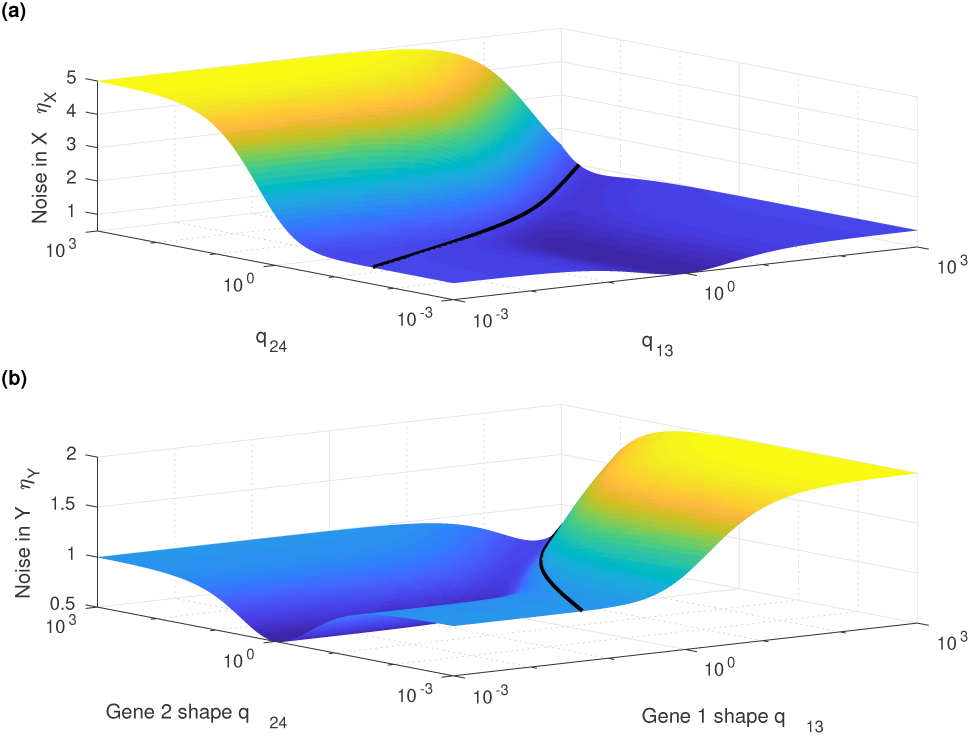
Noise in the RNA production in a head-to-head configuration as a function of their relative durations of closed and open complex formations. (A) gene 1 and (B) gene 2. The black curves denote unity, and *q*_12_=2.

**FIG. 4.**
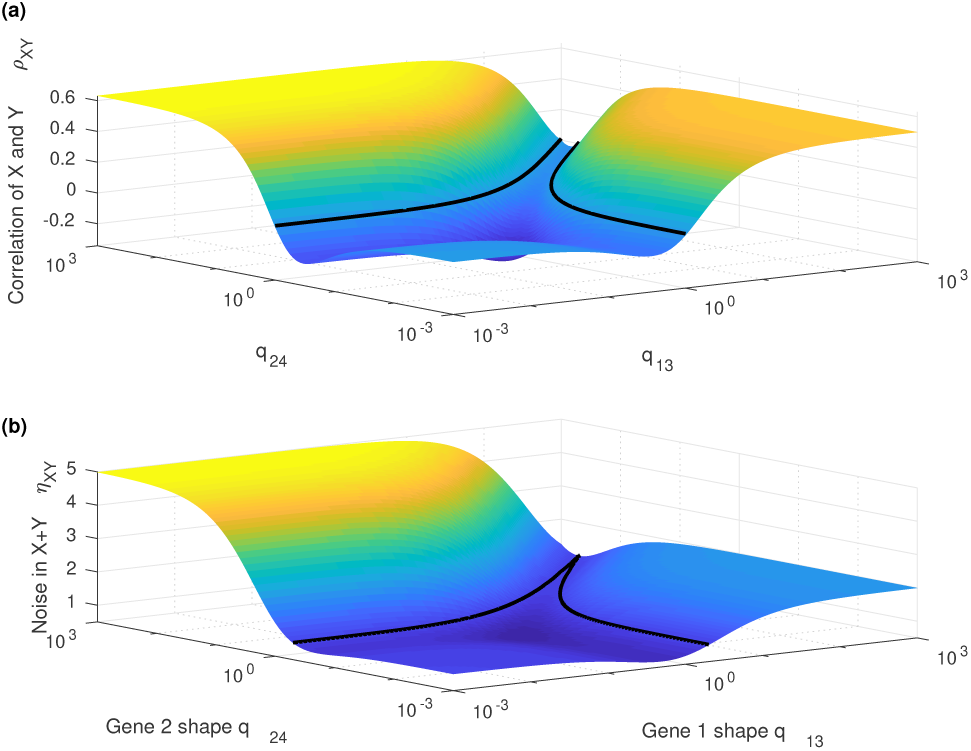
Correlation and total noise (tandem configuration) as a function of the relative durations of closed and open complex formation. (A) Correlation between the RNA production kinetics of two head-to-head genes. (B) Noise of RNA production of a gene with two initiation sites. The black curves denote zero or unity, and *q*_12_ = 2.

Region A: For *q*_24_ > 1, *q*_24_ > *q*_13_, the expression of gene 2 is most limited at the open complex formation, while that of gene 1 is more symmetric. As such, the promoter region is mostly occupied, and gene 1 must express either fast or rarely. In the former case, there is a burst of production of proteins *X* after each *Y*, so the expression of the two genes is positively correlated, and while gene 2 is Poissonian, solely controlled by its open complex formation process, gene 1 is highly noisy as the geometric burst of RNA is separated by the gaps created by the other gene. In the latter case, the expression of gene 1 is controlled by uniform random productions and the correlation vanishes. Specifically, in the latter case, the noise of gene 1 is 1 + 2 *q*_12_ (super-Poissonian) and gene 2 is Poissonian. The correlation for large *q*_12_ is 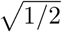, which is maximal for the configuration, while for small *q*_12_ the correlation vanishes. The part *q*_13_ > *q*_24_, *q*_13_ > 1 is symmetric. Note that the bias *q*_12_ controls the upper bounds for noise and correlation.

Region B: Here, both genes are limited at the closed complex formation. Thus, the promoter region is rarely occupied, as the expression is limited by an RNAP finding the gene and initiating transcription. This causes the expression of both genes to be Poissonian, as each is limited by a single step, and uncorrelated, as their activities do not interfere at the promoter region.

Region C: Both genes are limited at the open complex formation, which makes them to alternate in occupying the shared promoter region. The noise is set by the bias *q*_12_, which determines the gene more disturbed by the activity of the other. Specifically, the noise of gene 1 equals 1*/*2+*q*_12_ */* 2 and the noise of gene 2 equals 1*/*2+ *q*_12_^−1^ */* 2. As the genes inhibit each other by competing for the shared promoter region, the expression patterns are anticorrelated.

Region D: For *q*_13_ ∼ 1, *q*_24_ < 1, gene 2 is limited during the closed complex formation, so it does not block the shared promoter area. Meanwhile, gene 1 is limited at both stages, making its RNA production to be sub-Poissonian. The expression of gene 2, originally Poissonian, becomes affected by periods of inactivity as gene 1 employs the promoter, increasing the noise, as controlled by the bias, yielding noise of 1+*q*_12_^−1^ */* 2. The correlation is negative, as gene 1 inhibits the expression of gene 2. The part *q*_13_ < 1, *q*_24_ ∼ 1 is symmetric.

Region E: Both genes have similar closed and open complex formation durations, resulting in low noise in a non-bidirectional configuration. If their closed complex formation durations are similar (i.e. *q*_12_ ∼ 1), both genes are of low noise (∼ 7*/*9) and their expression is anticorrelated (∼−2*/*7), as they alternate in activity. Otherwise, one is of low noise (∼ 5*/*9), unaffected by the configuration, while the other is of high noise, with its expression being disturbed by the frequent gaps caused by the other. Specifically, the noise is 5*/*9 +2 *q*_12_ */* 9 for gene 1 and 5*/*9+2 *q*_12_^−1^ */* 9 for gene 2. The correlation is negative, with a maximum of −2*/*7 at *q*_12_ = 1, and minima of 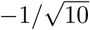 at *q*_12_ → 0 and *q*_12_ → ∞.

In summary, for coupled gene activity, one (or both) genes must not be limited at the closed complex formation alone. When coupled, both genes are low noise only if both feature similar relative closed-to-open complex durations. In this case, their expression is likely anti-correlated. If the relative closed-to-open complex durations differ, one is of high noise and the other of low noise, while their expression is, surprisingly, positively correlated. While our analysis lacks processes other than transcription initiation, the presence of e.g. first-order degradation pulls the noise toward unity and the correlation toward zero, leaving the conclusions qualitative useful.

### C. Noise in a gene with two initiation sites: model predictions and empirical validation

Next, we investigate the dynamics of a gene controlled by a promoter with two TSSs (cf. Fig S1C). This is common in *E. coli* [39] and more so in, e.g. plant mitochondria [40]. The configuration is readily accommodated by our model, by considering the dynamics of *X* + *Y*. As the mean and variance of *X* + *Y* follow the mean and covariance of (*X, Y*), the results can be derived from those obtained in the previous section.

Fig 4B shows the noise for *X* + *Y*, representing the RNAs produced through either TSS. The noise is low only if both TSSs exhibit production dynamics with low noise, i.e. in the regions C, D, or E. Compared to individual TSSs, the RNA number fluctuations are lower, being suppressed by the negative correlation. If one TSS exhibits highly noisy production (region A), the RNA numbers become highly noisy, regardless of the dynamics of the other TSS. Finally, in region B, the production is exponential-like, as multiple TSSs only increase the RNAP to promoter binding affinity, which makes their dynamics indiscernible from that of a single TSS.

To validate our predictions, we observed transcription in live *E. coli* at the single-RNA level in various constructs. Three of the constructs feature synthetic genes whose production is controlled by a single promoter (specifically P_lacO3O1_, P_tetA_, and P_BAD_ (cf. Fig S1A). The other constructs feature pairs of genes sharing promoter elements. One of these constructs is P_lacO3O1-lacO3O1_, with overlapping lacO3O1 promoters in the opposite strands (cf. Fig S1B), with the reporter being on a single side. In the other two such constructs, the expression is controlled by a P_lacO3O1-tetA_ or a P_lacO3O1-BAD_ dual-tandem promoters (cf. Fig S1C). In all these, the expression of the lacO3O1 promoter is modulated by the IPTG concentration, an inducer for the lac promoter[41]. Meanwhile, aTc concentration is held constant at 15 ng/ml, in order to trigger full expression of the tetA promoter. Similarly, L-arabinose concentration is held constant at 0.1%, to trigger full expression of the BAD promoter. In all cases, RNA production dynamics was measured by time-lapse microscopy imaging using MS2-GFP tagging (Methods).

Using our models, we aim to predict the behavior of the pairs of genes sharing promoter elements, given knowledge of the behavior of the constituent genes when not sharing such elements. I.e., we test whether, from the measured dynamics of RNA production of P_lacO3O1_, P_tetA_ [17], and P_BAD_, one can predict the kinetics of P_lacO3O1-lacO3O1_, P_lacO3O1-tetA_, and P_lacO3O1-BAD_.

For this, we first extracted the number of RNAs in each cell in the first and the last frame of the time series for all the constructs in each condition. These data were used to estimate the mean and standard deviation the production intervals, and the most likely (maximum likelihood fit) model of Eq (S1) for the single promoters. The estimated intervals are shown in Table II, along with the model parameters of Eq (S1) where applicable. A Wald test testing for a specific mean and standard deviation was used to compute a p-value to confirm that the model predicts the mean and variance of the RNA distributions.

**TABLE II.** Estimated RNA production intervals for each of the promoter constructs. The table shows the promoter, induction, estimated paramater of model Eq (S1) for the single promoters, the estimated mean, standard deviation (sd), and noise (coefficient of variation) of the RNA production intervals, and the p-value of the test of model versus data.

| Promoter | IPTG ($\mu\text{M}$) | $k_1^{-1}, k_2^{-1}$ (s) | $k_3^{-1}, k_4^{-1}$ (s) | Mean (s) | Sd (s) | Noise | P-value |
| --- | --- | --- | --- | --- | --- | --- | --- |
| lacO3O1 | 1000 | 362.3 | 737.3 | 1099.6 | 821.5 | 0.558 | 0.446 |
| lacO3O1 | 5 | 25.8 | 1236.8 | 1262.6 | 1237.0 | 0.960 | 0.273 |
| tetA | - | 287.4 | 385.5 | 672.9 | 480.9 | 0.511 | 0.075 |
| BAD | - | 1036.7 | 333.7 | 1370.4 | 1089.1 | 0.632 | 0.059 |
| lacO3O1-tetA | 1000 | - | - | 702.2 | 638.8 | 0.828 | 0.604 |
| lacO3O1-tetA | 5 | - | - | 1111.6 | 1089.1 | 0.960 | 0.164 |
| lacO3O1-lacO3O1 | 1000 | - | - | 1659.3 | 1437.0 | 0.750 | 0.112 |
| lacO3O1-lacO3O1 | 5 | - | - | 2205.9 | 2119.3 | 0.923 | 0.971 |
| lacO3O1-BAD | 1000 | - | - | 866.8 | 612.9 | 0.500 | 0.099 |
| lacO3O1-BAD | 5 | - | - | 1274.7 | 1248.9 | 0.960 | 0.698 |

The results in Table II indicate that changing IPTG concentration alters the noise of the lacO3O1 promoter in addition to changing its mean expression rate, which is expected to be due to changes in the open-to-closed complex duration ratio, in agreement with previous reports [19]. The p-values indicate that there is no evidence that any of the models fit the measurements poorly. We also extracted the intervals from the full time series for several of the cases (about 120 frames, one every second) to verify that they are correctly estimated (see Table S-II).

Next, using the above parameters, we constructed the models for the dual promoters. The obtained models are shown in Table III. The mean and standard deviation show an agreement with the empirical data, while the noise and correlation indicate that that promoters operate at different regions of the open-to-closed complex ratio space. The results indicate that the model predicts the behavior of the dual-promoter measurements well, and that the noise is modulated by the change in the coordination between the two promoters in the dual promoter construct.

**TABLE III.** Models derived for the dual promoters from the individual promoter fits of Table II using the model with interactions during transcription initiation. The table shows the promoter/induction scheme, the mean and standard deviation (sd) of the RNA production intervals and the correlation between the RNA numbers assuming the derived models, and the p-value of the test of model versus data.

| Promoter | IPTG ($\mu\text{M}$) | Mean (s) | Sd (s) | Noise | Correlation | P-value |
| --- | --- | --- | --- | --- | --- | --- |
| lacO3O1-lacO3O1 | 1000 | 1836.9 | 1685.2 | 0.842 | -0.188 | 0.168 |
| lacO3O1-lacO3O1 | 5 | 2499.3 | 2486.5 | 0.990 | -0.010 | 0.931 |
| lacO3O1-tetA | 1000 | 701.4 | 616.1 | 0.772 | -0.166 | 0.603 |
| lacO3O1-tetA | 5 | 1190.4 | 1212.9 | 1.038 | +0.172 | 0.123 |
| lacO3O1-BAD | 1000 | 901.3 | 731.4 | 0.659 | -0.070 | 0.141 |
| lacO3O1-BAD | 5 | 1240.0 | 1230.9 | 0.985 | +0.105 | 0.620 |

As our methodology cannot identify which of the steps correspond to *k*_1_^−1^ and *k*_3_^−1^ in Table II, we also considered the alternative step ordering. The dual-promoter model fits had a p-values < 3.964 × 10^−3^ in for the lacO3O1-tetA construct at 5 μM IPTG, and p-values < 4.614×10^−3^ for the lacO3O1-BAD construct at 1000 μM IPTG, indicating that the alternatives are not likely for lacO3O1 at 5 μM and BAD. The step order for lacO3O1 at 1000 μM IPTG and tetA cannot be resolved from these data, but the alternatives result in a qualitatively similar dual-promoter models and p-values > 0.116. For the constructs containing these two promoters, we report the most likely models, all suggesting the order specified in Table II. These findings are also supported by prior evidence using a different methodology[19].

The fact that the measurements fall into the different regions of operation (see Fig 4 and Table I) is apparent in Fig S2, Fig S3, and Fig S4. Namely, the high IPTG condition falls into region E for the lacO3O1-lacO3O1 and lacO3O1-tetA, and into region D for the lacO3O1-BAD construct. At low IPTG, the lacO3O1-lacO3O1 transits into region C, as both promoters are modulated by the changes in the inducer concentration, while the lacO3O1-tetA and lacO3O1-BAD transit into (opposite directions) of region A. This explains the widely different noise levels in the measured intervals, which are well predicted by our models in each case.

Finally, we attempted to predict the mean, noise, and intervals in a dual-promoter measurement assuming that there were no interactions between the two promoters. The results in Table IV show that the associated model fails to explain the observed dual-promoter behavior. Note that the models are also unaffected by the (*k*_1_, *k*_3_) identifiability problem. While the mean and noise of the system consisting of two independent promoters trivially follow from their independent components, the time intervals of the combined production do not. In particular, the intervals are not independent. We also considered the possibility that while the promoters might have interactions, their expression levels may be altered by the other promoter utilizing the same finite pool of RNA polymerases. For this, we assumed that the number of RNA polymerases modulate the closed complex formation rate (i.e. *k*_1_ = *R* 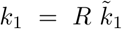 where 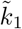 is the per-polymerase closed complex formation rate, and *R* represents an RNA polymerase), which will cause a slight reduction of the closed complex formation rate, as determined by the closed to open-to-closed complex duration ratio of the other promoter. Any of these models (all *R* and all step orders) failed to explain the behavior of our dual-promoter measurements as well. The effects are most extreme for *R* = 2, but we verified that models for other *R* have no better fit. Our model is recovered at *R* = 1 and the independent model without polymerases is recovered at *R* = ∞.

**TABLE IV.**
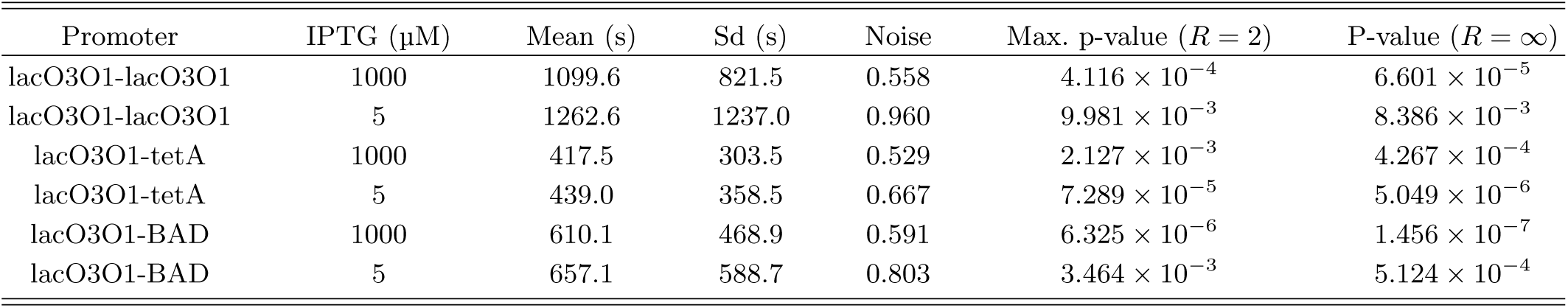
Null models derived for the dual independent promoters from the individual promoter fits of Table II. The table shows the promoter/induction scheme, and the mean and standard deviation (sd) of the RNA production intervals assuming the null models, and the p-value of the test of model versus data for maximally (*R* = 2) and minimally (*R* =∞) RNA polymerase starved null models.

| Promoter | IPTG (μM) | Mean (s) | Sd (s) | Noise | Max. p-value ($R = 2$) | P-value ($R = \infty$) |
| --- | --- | --- | --- | --- | --- | --- |
| lacO3O1-lacO3O1 | 1000 | 1099.6 | 821.5 | 0.558 | $4.116 \times 10^{-4}$ | $6.601 \times 10^{-5}$ |
| lacO3O1-lacO3O1 | 5 | 1262.6 | 1237.0 | 0.960 | $9.981 \times 10^{-3}$ | $8.386 \times 10^{-3}$ |
| lacO3O1-tetA | 1000 | 417.5 | 303.5 | 0.529 | $2.127 \times 10^{-3}$ | $4.267 \times 10^{-4}$ |
| lacO3O1-tetA | 5 | 439.0 | 358.5 | 0.667 | $7.289 \times 10^{-5}$ | $5.049 \times 10^{-6}$ |
| lacO3O1-BAD | 1000 | 610.1 | 468.9 | 0.591 | $6.325 \times 10^{-6}$ | $1.456 \times 10^{-7}$ |
| lacO3O1-BAD | 5 | 657.1 | 588.7 | 0.803 | $3.464 \times 10^{-3}$ | $5.124 \times 10^{-4}$ |

We conclude that our model of closely-spaced promoters that assumes interactions between the promoters is the one that well predicts the measurements in each setting, for both the head-to-head and tandem constructs. Relevantly, our models reveal that the observed changes arise from changes in the coordination between the two coupled transcription start sites of our synthetic constructs.

## IV. CONCLUSION

We analyzed a stochastic model of two genes in a head-to-head configuration as a function of whether each gene is rate-limited during the closed and/or open complex formation. Compared to individual genes, in the bidirectional configuration, the transcription activity is slower and noisier in both genes, as each gene interferes with the activity of the other, allowing two genes with sub-Poissonian dynamics to exhibit super-Poissonian dynamics when coupled. Importantly, provided information on the kinetics of the constituent promoters when not sharing promoter elements, the models were shown to be able to predict well the behavior of the pairs of the same genes when sharing promoter elements, implying that they capture accurately the effects of the complex interference caused by the sharing of elements.

We found that for such prediction to be accurate, the models have to account for the two-rate-limiting step kinetics of active transcription in *E. coli*. In particular, the time-length of such rate limiting steps, namely the closed and open complex formations, controls not only the expression rate and noise of each gene (as in isolated genes, see e.g. [15]), but also the kinetics of the temporal gaps caused by the transcription events of the opposite gene. This programs the behavior intricately: a similar rate-limiting step structure combined with a rate-limiting open complex formation is required for both genes to feature low noise; otherwise one tends to be highly noisy. Also, orderly systems tend to exhibit strong negative correlation, while the genes alternate expression, but the correlation can be lost or become positive if the open-to-closed complex formation time-lengths are incompatible. As such, not only the mean and variance of the durations of each stage, but also the mechanistic underpinnings, affect the dynamics of closely-spaced gene-pairs, implying that promoters with seemingly identical dynamics in isolation may differ widely in their dynamics in a closely-spaced configuration. Relevantly, as shown, the results generalize to the behavior of individual genes with multiple transcription initiation sites.

Overall, these results suggest that, in *E. coli*, the kinetics of the rate-limiting steps in active transcription needs to be considered for dissecting the dynamics of pairs of genes sharing promoter elements. In this regard, we find it to be striking that pairs of closely-spaced promoters, by tuning the kinetics of their closed and open complex formation (which are sequence dependent and, thus, evolvable) tunes the orderliness of the whole genepair. This new knowledge provides an important route to follow in the engineering of pairs of closely-spaced promoters with desired dynamics and contributes to a better understanding of the dynamics of natural pairs of closely spaced genes and their potential role in the gene expression programs of *E. coli*.

## ACKNOWLEDGMENTS

The work was supported by the Alfred Kordelin Foundation (AH), the Vilho, Yrjo and Kalle Vaisala Foundation (SMDO), the Tampere University of Technology President’s Graduate Programme (RN-V), the Jane and Aatos Erkko Foundation (610536; ASR), and Academy of Finland (295027 and 305342; ASR). The funders had no role in study design, data collection and analysis, decision to publish, or preparation of the manuscript. The authors declare that they have no conflict of interest.

