## Supplementary material for "Transcription closed and open complex formation coordinate expression of genes with a shared promoter region"

### MODELS

#### Production intervals of an isolated single gene

For each individual gene, we assume the following model of RNA production [S1], whose dynamics are expected to follow stochastic chemical kinetics [S2, S3]:

$$P_0 \xrightarrow{k_1} I_1 \xrightarrow{k_3} P_0 + X \tag{S1}$$

where $P_0$ and $I_1$ represent unoccupied and occupied states of the promoter, respectively, while $X$ the messenger RNA produced by the gene.

The reactions with rates $k_1$ and $k_3$ represent the main rate limiting steps during transcription. Typically, in bacteria, transcription is rate-limited by events during transcription initiation [S1, S4]. The two most time-consuming events during transcription initiation are the closed complex formation, which involves $\sigma$-factor recruitment, diffusion along the DNA and specific binding of the RNA polymerase and the associated machinery at transcription start site; and the open complex formation, which involves unwinding and compaction of the DNA and assembly of the RNA polymerase clamp [S1, S4]. We assume that the cell is not starved of RNA polymerases, which is applicable under normal growth conditions, such that the effective polymerase abundance can be encoded in $k_1$. The steps and the model used here are illustrated in Fig. S1A.

Physical arguments suggest that the waiting time for a reaction with rate $k$ reaction ought to follow an exponential distribution with the corresponding rate [S5]. As the closed and open complex formation are complex processes, it is not obvious that they should feature such an elementary dynamical scheme. However, the model has been found to successfully capture the expression of several promoters in *E. coli* under various conditions when sampled at a 1 minute resolution [S6–S10].

Other events such as transcription elongation are expected to play a negligible role, as elongation takes tens of seconds [S11], while the RNA production intervals are in the order of hundreds of seconds [S6–S10]. Regardless, elongation does not affect the RNA production intervals on expectation, unless polymerase traffic forms due to a very fast rate of initiation [S12]. These assumptions can be expected to hold even under extreme conditions [S13].

Following the model of Eq. (S1), the time intervals between the production of consecutive RNAs ${\tau_X}^{(1)}$ are independent and identically distributed with the following density:

$$f_{{\tau_X}^{(1)}}(t) = \left(f_{k_1} * f_{k_3}\right)(t) \quad = \left(\tfrac{k_3}{k_3 - k_1}\right)\, f_{k_1}(t) + \left(\tfrac{k_1}{k_1 - k_3}\right)\, f_{k_3}(t) \tag{S2}$$

where $f_{k_i}(t) \doteq k_i\, \exp(-k_i\, t)$ is the exponential density of an individual reaction with a rate of $k_i$, and $f * g$ is the convolution of $f$ and $g$. The singularity $k_3 \to k_1$ is removable, not affecting the analysis [S14].

The moments can be extracted from the density:

$$\mathbb{E}\left[{{\tau_X}^{(1)}}^n\right] = \left(\tfrac{{k_1}^{n+1} - {k_3}^{n+1}}{k_1 - k_3}\right)\, \tfrac{n!}{(\, k_1\, k_3\, )^n} \tag{S3}$$

from which the mean and variance follow:

$$\begin{aligned} \mathbb{E}\left[{\tau_X}^{(1)}\right] &= {k_1}^{-1} + {k_3}^{-1} \\ \mathbb{V}\mathrm{ar}\left[{\tau_X}^{(1)}\right] &= {k_1}^{-2} + {k_3}^{-2} = \underbrace{\left(\frac{1 + {q_{13}}^2}{(1 + q_{13})^2}\right)}_{\eta_{X^{(1)}}}\ \underbrace{\mathbb{E}\left[{\tau_X}^{(1)}\right]^2}_{{\lambda_1}^2 \doteq {\mu_{X^{(1)}}}^{-2}} \end{aligned} \tag{S4}$$

where $q_{13} \doteq k_1 \,/\, k_3$ is the ratio between the durations of open and closed complex formation (${k_3}^{-1}$ and ${k_1}^{-1}$, respectively). The mean and noise (squared coefficient of variation, variance to mean-squared ratio; see the next section why this is the preferred definition of noise) can be freely varied, noise being determined by the open-to-closed complex formation duration ratio. The noise is unity for $q_{13} \to 0$ and $q_{13} \to \infty$ (maximum), for a single rate limiting step, and one half at $q_{13} = 1$ (minimum) for two rate limiting steps of equal duration. The parameterization $\lambda_1$, $q_{13}$ represents the decomposition of the mean and variance to the independent terms.

In this work, we use the properties derived by analytical means, but the results can be verified using Monte Carlo simulations [S3], as in Fig. 2B and Fig. 2C.

**Relationship between production intervals and RNA numbers**

While the distribution of transcription intervals best characterizes the dynamical behavior of the transcription process, perhaps a more meaningful measure from the point of view of cell faith or cell population is the resulting distribution of RNA (or rather protein) numbers. This stems from the fact that cellular decisions are carried out by proteins, which, in bacteria, tend to follow the RNA numbers [S15].

For independent production intervals $\tau_Z$, the distribution of produced RNA numbers $Z(t)$ at time $t$ has the complementary cumulative density:

$$\mathbb{P}[\, Z(t) \geq z \,] = \mathbb{P}\left[ \sum_{i=1}^{z} \tau_{Z_i} \leq t \right] = (\, H * {f_{\tau_Z}}^{*\,z} \,)\,(t) \tag{S5}$$

where $\tau_{Z_i}$ is the $i$:th production interval, $f_{\tau_Z}(t)$ their density, $H(t) \doteq \mathbb{I}\{\, t \geq 0 \,\}$ is the unit step function, and $f^{*\,k}$ is the $k$:th convolution power of $f$. This assumes that there are zero RNAs $Z(0) = 0$ at time $t = 0$, and that the process starts exactly at $t = 0$. However, for long-term behavior ($t \to \infty$) the effects of the initial conditions vanish. That is, the two are intimately linked as suggested above, disregarding knowledge of the previous state.

It can be shown that in the long-term ($t \to \infty$), the mean and variance of produced RNA per unit time are [S16]:

$$\begin{aligned} \mu_Z &\doteq \lim_{t\to\infty} \frac{\mathbb{E}[\, Z(t) \,]}{t} = \mathbb{E}[\tau_Z]^{-1} \\ \eta_Z\, \mu_Z &\doteq \lim_{t\to\infty} \frac{\mathbb{V}\mathrm{ar}[\, Z(t) \,]}{t} = \mathbb{V}\mathrm{ar}[\tau_Z]\ \mathbb{E}[\tau_Z]^{-3} \end{aligned} \tag{S6}$$

i.e. the mean number of RNAs produced per unit time ($\mu_Z$) equals the inverse interval mean, while the Fano factor (variance over the mean) of the RNA numbers ($\eta_Z$) equals the squared coefficient of variation of the production intervals.

If transcription is an elementary chemical process, it follows that the production intervals are exponentially distributed [S5], and the produced RNA numbers follow a Poisson distribution with the appropriate rate. Regardless of the production rate $\mu_Z$, the Fano factor of the RNA numbers (which equals the squared coefficient of variation of the transcription intervals) in such process is constant, $\eta_Z = 1$. The dynamics of more complex processes can be characterized independently of their mean using this statistic for the relative fluctuations ("noise"): processes that are more (less) stochastic than a Poisson process with the corresponding rate, are called super-Poissonian (sub-Poissonian).

In live cells, the active RNA numbers are determined not only by transcription, but also by RNA degradation and dilution of RNA through cell divisions. Each process affects both the mean and stochasticity of the active RNA numbers, and thus the protein numbers [S17, S18]. However, as their contribution can be approximately decomposed into (following the approximations from [S17]):

$$\eta_{\text{ active }} \approx \left(\frac{\bar{k}/2}{\bar{k}+\bar{d}/2}\right) \eta_{\text{ production }} + \left(1 - \frac{\bar{k}/2}{\bar{k}+\bar{d}/2}\right) \eta_{\text{ decay}} \tag{S7}$$

where $\bar{k}$ and $\bar{d}$ are the mean transcription and decay (degradation plus dilution) rates, respectively, the mean and noise of produced RNA numbers well characterizes the noise contributed by the transcription process to the diversity of cellular phenotypes. For first-order-like degradation and dilution, any changes in the noise introduced by transcription will still be reflected in the RNA numbers, but the effect can be made less prominent by the other processes, as the above suggests. More complex RNA degradation and/or partitioning at cell division are known to result in changes of the autocorrelation decay (i.e. short-term coordination) of the RNA numbers, but are not expected to affect the applicability of the above scheme [S17].

**Production intervals for genes with a shared promoter**

Next, we analyzed the RNA production intervals under the assumption that two genes, arranged in a head-to-head configuration, share interactions during their transcription initiation. We assume that the promoter can be in three different states: free, or occupied for transcription initiation into either of the two directions (see Fig. 2A and Fig. S1B). The closed complex formation, which includes the assembly of the RNA polymerase holoenzyme and DNA binding [S1, S4], can occur in parallel in our model, while the open complex formation cannot occur simultaneously for expressing the two genes, as the shared promoter region can be only employed for expression into one of the directions. This model has been employed in previous literature [S19].

The model for the head-to-head configuration with a shared promoter is:

$$\begin{aligned} P_0 &\xrightarrow{k_1} I_1 \xrightarrow{k_3} P_0 + X \\ P_0 &\xrightarrow{k_2} I_2 \xrightarrow{k_4} P_0 + Y \end{aligned} \tag{S8}$$

where $P_0$, $I_1$, and $I_2$ represent the different states of the promoter, and $X$ and $Y$ the messenger RNAs of the two genes. Here, $k_1$ and $k_2$ are the closed complex formation rates of the two genes, and $k_3$ and $k_4$ the corresponding open complex formation rates.

From the perspective of producing only $X$, the model can be written as:

$$I_2 \underset{k_2}{\overset{k_4}{\rightleftharpoons}} P_0 \xrightarrow{k_1} I_1 \xrightarrow{k_3} P_0 + X \tag{S9}$$

Following these reactions, the production intervals of $X$ are independent and identically distributed according to the density:

$$f_{\tau_X}(t) = \left( \sum_{k=0}^{\infty} \left( \tfrac{k_2}{k_1+k_2} \right)^k f_{k_1+k_2}{}^{*k} * f_{k_4}{}^{*k} * \left( \tfrac{k_1}{k_1+k_2} \right) f_{k_1+k_2} * f_{k_3} \right)(t) \tag{S10}$$

where the sum runs over all possible number of visits through $I_2 = 1$ (i.e. intermittent initiations to express gene 2) prior to committing to the expression of gene 1. A more detailed treatment can be found in [S14]. Again, the mean and variance are readily extracted from the density:

$$\begin{aligned} \mathbb{E}[\tau_X] &= \left(1 + \tfrac{k_2}{k_4}\right) k_1{}^{-1} + k_3{}^{-1} \\ \mathbb{V}\mathrm{ar}[\tau_X] &= \left( \left(1 + \tfrac{k_2}{k_4}\right)^2 + 2\,\tfrac{k_1}{k_4}\,\tfrac{k_2}{k_4} \right) k_1{}^{-2} + k_3{}^{-2} \end{aligned} \tag{S11}$$

which emphasizes how the mean and variance of the closed complex formation are modulated by the expression of gene 2 (cf. Eq. (S4)), while those of the open complex formation are unaffected. Note that in this model, $\tau_X$ and the similar $\tau_Y$, which follows from the symmetry, are not independent.

For the purposes of determining the dependence between the two promoters, we also determined how the model parameters affect the dynamics of $X + Y$, that is, the total number of RNAs produced by either gene. Again, the density for the intervals can be written out as:

$$f_{\tau_{X+Y}}(t) = \left( \left( \tfrac{k_1}{k_1+k_2} \right) f_{k_1+k_2} * f_{k_3} + \left( \tfrac{k_2}{k_1+k_2} \right) f_{k_1+k_2} * f_{k_4} \right)(t) \tag{S12}$$

where the two terms represent the two paths to produce an RNA in the model. This density results in mean and variance of:

$$\begin{aligned} \mathbb{E}[\tau_{X+Y}] &= \left(1 + \tfrac{k_1}{k_3} + \tfrac{k_2}{k_4}\right) (k_1 + k_2)^{-1} \\ \mathbb{V}\mathrm{ar}[\tau_{X+Y}] &= \left(1 + \left(\tfrac{k_1}{k_3} - \tfrac{k_2}{k_4}\right)^2 + 2\,\tfrac{k_1}{k_3}\,\tfrac{k_2}{k_3} + 2\,\tfrac{k_1}{k_4}\,\tfrac{k_2}{k_4}\right) (k_1 + k_2)^{-2} \end{aligned} \tag{S13}$$

TABLE S-I. Approximate expressions for the noise and correlation of the two genes in a head-to-head configuration. The expressions are approximations of the expressions shown in Eq. (S14) and Eq. (S15) about the ideal points of the regions.

| Region | Condition | Noise $\eta_X$ | Noise $\eta_Y$ | Correlation $\rho_{XY}$ |
|---|---|---|---|---|
| A | $q_{24} > 1,\, q_{24} > q_{13}$ | $1 + 2\,q_{12}$ | $1$ | $+\sqrt{2 + {q_{12}}^{-1}}^{\,-1}$ |
| A | $q_{13} > q_{24},\, q_{13} > 1$ | $1$ | $1 + 2\,{q_{12}}^{-1}$ | $+\sqrt{2 + q_{12}}^{\,-1}$ |
| B | $q_{13},\, q_{24} < 1$ | $1 - 2\,q_{13}$ | $1 - 2\,q_{24}$ | $-\left(\sqrt{{q_{12}}^{-1}}\,q_{13} + \sqrt{q_{12}}\,q_{24}\right)$ |
| C | $q_{13} \sim q_{24} > 1$ | $1/2 + q_{12}/2$ | $1/2 + {q_{12}}^{-1}/2$ | $-{q_{13}}^{-1}/2 \;\sim\; -{q_{24}}^{-1}/2$ |
| D | $q_{13} \sim 1,\, q_{24} < 1$ | $1/2$ | $1 + {q_{12}}^{-1}/2$ | $-\sqrt{1 + q_{12}}\;q_{24}/2$ |
| D | $q_{13} < 1,\, q_{24} \sim 1$ | $1 + q_{12}/2$ | $1/2$ | $-\sqrt{1 + q_{13}}\;q_{13}/2$ |
| E | $q_{13} \sim q_{24} \sim 1$ | $5/9 + 2\,q_{12}/9$ | $5/9 + 2\,{q_{12}}^{-1}/9$ | $-\left(1 + q_{12}\right)/\sqrt{2 + 5\,q_{12}}/\sqrt{5 + 2\,q_{12}}$ |

The intervals $\tau_{X+Y}$ can also be interpreted in another sense. Namely, if a gene features a pair of transcription start sites, each of which can recruit the transcriptional machinery, but share interactions at the promoter region (see Fig. S1C), this quantity represents the number of RNAs produced through initiations through either of the sites. In such case, $X$ and $Y$ cannot be separately measured for practical purposes, as the number of produced RNA corresponds to $X + Y$, which is also what is relevant for the cellular phenotype.

**Effects of shared promoter area on RNA numbers**

The effects of transcriptional dynamics on the long-term RNA numbers can be determined from Eq. (S11) using Eq. (S6). The noise for the RNA $X$ in the head-to-head two gene configuration is:

$$\eta_X = \mathbb{V}\mathrm{ar}[\tau_X]\;\mathbb{E}[\tau_X]^2 = \frac{\left(1 + q_{24}\right)^2 + 2\;q_{12}\;{q_{24}}^2 + {q_{13}}^2}{\left(\,1 + q_{13} + q_{24}\,\right)^2} \tag{S14}$$

where $q_{ij} \doteq k_i/k_j$. Again, $\eta_Y$ follows from the symmetry. By itself, this expression might not be particularly insightful, but can be visualized (see Fig. 2). In addition, the expression can be simplified in the different regions of the parameter space, as discussed in the main manuscript and tabulated in Table S-I.

The correlation between the RNA numbers produced by the two genes can be derived from the marginal statistics $\mathbb{E}[X]$, $\mathbb{V}\mathrm{ar}[X]$, $\mathbb{E}[Y]$, and $\mathbb{V}\mathrm{ar}[Y]$ and the sum statistics $\mathbb{E}[X+Y]$ and $\mathbb{V}\mathrm{ar}[X+Y]$. Namely, the correlation can be found through the basic probability identities of variance:

$$\rho_{XY} \doteq \mathbb{C}\mathrm{orr}[X,Y] = \frac{\mathbb{C}\mathrm{ov}[X,Y]}{\sqrt{\mathbb{V}\mathrm{ar}[X]\,\mathbb{V}\mathrm{ar}[Y]}} = \frac{1}{2}\,\frac{\mathbb{V}\mathrm{ar}[X+Y] - \mathbb{V}\mathrm{ar}[X] - \mathbb{V}\mathrm{ar}[Y]}{\sqrt{\mathbb{V}\mathrm{ar}[X]\,\mathbb{V}\mathrm{ar}[Y]}} \tag{S15}$$

where $\rho_{XY}$ represents the Pearson's correlation coefficient between the RNA numbers produced by the two genes.

Specifically, in terms of $q_{12}$, $q_{13}$, and $q_{24}$, the correlation can be written as:

$$\rho_{XY} = -\frac{\sqrt{{q_{12}}^{-1}}\;q_{13}\;(1 - q_{13} + q_{24}) + \sqrt{q_{12}}\;q_{24}\;(1 + q_{13} - q_{24})}{\sqrt{(1+q_{24})^2 + 2\,q_{12}\,{q_{24}}^2 + {q_{13}}^2}\;\sqrt{(1+q_{13})^2 + 2\,{q_{12}}^{-1}\,{q_{13}}^2 + {q_{24}}^2}} \tag{S16}$$

which is shown in Fig. 4A. The expressions for the asymptotic correlation in the different regions discussed in the main manuscript are tabulated in Table S-I.

For a gene with multiple transcription start sites (cf. Fig. S1C), the correlation does not have a physical (measurable) meaning as the RNA products are indistinguishable, but in the model it quantifies the degree of cooperation between the two start sites, analogous to the head-to-head two gene configuration. The noise for this system is:

$$\eta_{X+Y} = \mathbb{V}\mathrm{ar}[\tau_{X+Y}]\;\mathbb{E}[\tau_{X+Y}]^{-2} = \frac{1 + 2\,{q_{12}}^{-1}\,{q_{13}}^2 + 2\,q_{12}\,{q_{24}}^2 + (q_{13} - q_{24})^2}{(\,1 + q_{13} + q_{24}\,)^2} \tag{S17}$$

which is shown in Fig. 4B. The asymptotic expressions for the different regions can also be derived (cf. Table S-I). In regions A and C, this follows the noisier gene, in region B the less noisy, in region D it interpolates between the two, and in region E it can even suppress the noise with respect to the marginals (i.e. $\eta_{X+Y} < \eta_X, \eta_Y$) if the promoters have similar expression rate (i.e. $q_{12} \sim 1 \Rightarrow \eta_X \sim \eta_Y \sim 7/9, \eta_{X+Y} \sim 5/9$).

**Production intervals of independent promoters**

The production dynamics of two independent promoters producing the same product can be derived from the dynamics of the individual promoters through Eq. (S6). We use this model as a null model to show that interactions between the promoters are needed to explain the measurement data.

Letting $\tau_X$ and $\tau_Y$ represent the production intervals for the marginals, the moments for the production interval $\mathring{\tau}_{X+Y}$ from either promoter of the system are:

$$\begin{aligned}\mathbb{E}\left[\mathring{\tau}_{X+Y}\right] &= \left(\mathbb{E}[\tau_X]^{-1} + \mathbb{E}[\tau_Y]^{-1}\right)^{-1} \\ \mathbb{V}\mathrm{ar}\left[\mathring{\tau}_{X+Y}\right] &= \left(\mathbb{V}\mathrm{ar}[\tau_X]\ \mathbb{E}[\tau_X]^{-3} + \mathbb{V}\mathrm{ar}[\tau_Y]\ \mathbb{E}[\tau_Y]^{-3}\right)\ \mathbb{E}\left[\mathring{\tau}_{X+Y}\right]^3\end{aligned} \tag{S18}$$

Further, we consider the possibility that the gene expression dynamics are perturbed due to the two promoters competing of a finite pool of RNA polymerases. Consider a single gene expression model of:

$$P_0 + R \xrightarrow{\tilde{k}_1} I_1 \xrightarrow{k_3} P_0 + R + X \tag{S19}$$

where $R$ represents the finite number of RNA polymerases at time $t$, $R_0$ being the initial number, and $\tilde{k}_1 \doteq k_1\, {R_0}^{-1}$ is the per-polymerase closed complex formation rate. If the other independent promoter perturbs the expression by reserving an RNA polymerase, the effective closed complex formation duration is:

$$\overset{*}{k}_1{}^{-1} \approx (1-p_2)\ \left(R_0\, k_1\, {R_0}^{-1}\right) + p_2\ \left((R_0-1)\, k_1\, {R_0}^{-1}\right) = \left(1 - p_2\, {R_0}^{-1}\right)\ k_1 \tag{S20}$$

where the probability of gene 2 being active is:

$$p_2 \doteq {k_4}^{-1} \,/\, \left(\overset{*}{k}_2{}^{-1} + {k_4}^{-1}\right) \tag{S21}$$

and $\overset{*}{k}_2$ and $p_1$ being defined in a symmetric fashion. Now, $\overset{*}{k}_1$ and $\overset{*}{k}_2$ can be solved from these recurrences and used as perturbed coefficients for the model with independent promoters to account for the finite RNA polymerase population shared by the two promoters. (These equations specify a quadratic system in $(\overset{*}{k}_1, \overset{*}{k}_2)$ which can be solved analytically. The other root is always negative, and can be disregarded when rates are concerned.)

**Model fitting**

Given a set of measured production intervals (or their bounds), the parameters of our models can be estimated in maximum likelihood sense using the densities of Eq. (S2) and Eq. (S12). The details are discussed e.g. in our previous work [S14]. An expectation maximization algorithm [S20] is possible, which lacks the problems that are associated with more general optimization procedures. Alternatively, the mean and variance of the production intervals can be inferred from the RNA distributions in each cell in the beginning and end of the time series through Eq. (S6), provided that the RNAs are virtually free of degradation, as is the case with MS2-GFP-tagged RNAs [S21]. The latter strategy was used to construct the models, as it facilitates profiling multiple constructs at a higher throughput, but we verified that the two approaches are comparable using the lacO3O1, tetA, and lacO3O1-tetA constructs (see Table S-II and the discussion that follows).

However, there is an inherent problem when doing inference with the model [S14]. Namely, equal (in the sense of probability distribution) RNA production intervals are produced by $(k_1, k_3) = (\lambda_-, \lambda_+)$ and $(k_3, k_1) = (\lambda_-, \lambda_+)$, which implies that while the possible values of $k_1$ and $k_3$ can be determined, it cannot be identified which is which. Typically, this conflict is resolved using further information [S8]. Here, we report the most likely case as determined by the model fits to the measurements in all conditions (the details are discussed below). The alternative cases for which we had no evidence to rule out were found not to differ significantly.

**Measured versus estimated time intervals**

To verify that the time intervals are accurately estimated from the RNA distributions of the first and last frame only, we extracted time intervals between production of consecutive RNAs from the full time series in three different constructs: lacO3O1, tetA, or a dual lacO3O1-tetA promoter. We further performed a Kolmogorov-Smirnov (K-S) test using the null hypothesis that the interval data are generated from the model distribution with mean and standard deviation equal to those estimated, in order to evaluate whether the two methods agree. The measurements of lacO3O1 and lacO3O1-tetA promoters were conducted as described in the previous sections, while the data from [S6] was used for the tetA promoter.

The results suggest that the two methods agree for the purposes of this study.

TABLE S-II. Measured RNA production intervals for lacO3O1, tetA, and lacO3O1-tetA constructs. The table shows the promoter, induction scheme, and the number, sample mean, standard deviation (sd), and noise (coefficient of variation) of the measured or estimated RNA production intervals, and the p-value for the interval data to differ from the expected distribution estimated from the first and last frame RNA distributions.

| Promoter | IPTG (µM) | Intervals | Mean (s) | Sd (s) | Noise | P-value |
|---|---|---|---|---|---|---|
| lacO3O1 | 1000 | 41 | 1096.1 | 849.7 | 0.601 | 0.301 |
| lacO3O1 | 5 | 35 | 1215.4 | 1229.9 | 1.024 | 0.438 |
| tetA | - | 254 | 617.0 | 367.4 | 0.355 | 0.066 |
| lacO3O1-tetA | 1000 | 39 | 703.1 | 649.0 | 0.852 | 0.230 |
| lacO3O1-tetA | 5 | 38 | 1277.4 | 1362.9 | 1.138 | 0.554 |

### SUPPLEMENTARY FIGURES

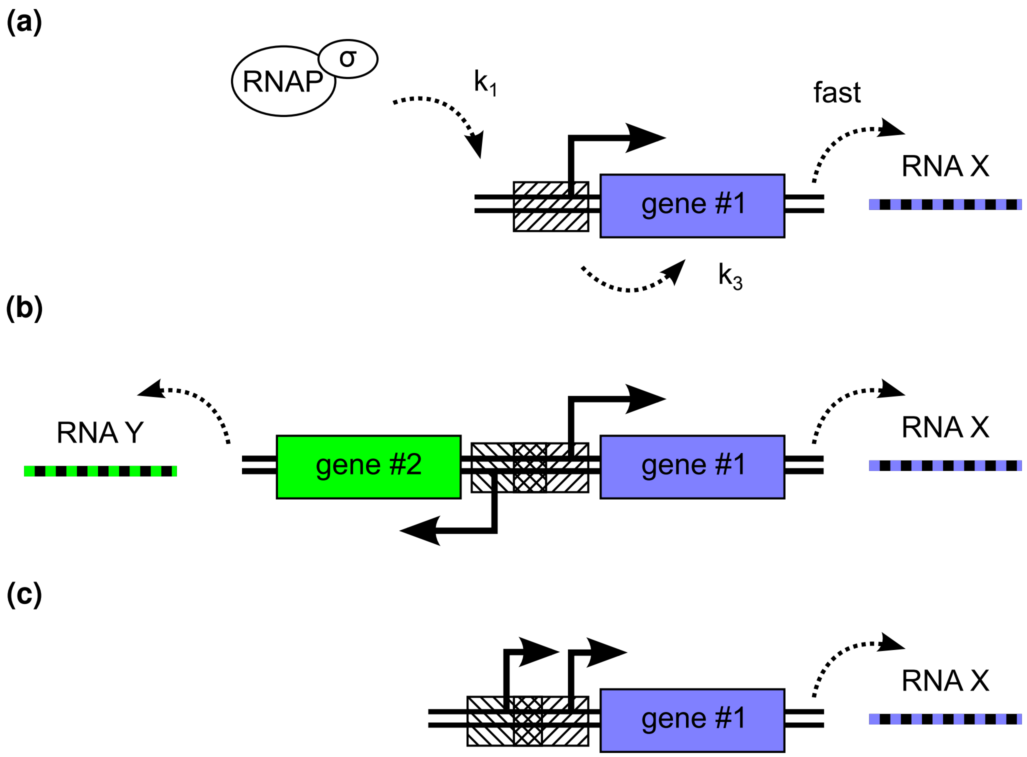


FIG. S1. Schematic of the modeled promoter architectures. (A) Schematic of the assumed transcription process of a single gene. The closed complex formation ($k_1$) consists of the RNA polymerase holoenzyme assembly and its binding to the promoter area (striped box). Open complex formation ($k_3$) follows, freeing the promoter. Subsequent steps such as elongation are expected to be fast compared to the closed and open complex formation. (B) Schematic of two genes in a head-to-head configuration with a shared promoter area (represented by the overlapping striped boxes) (C) Schematic of a gene with two transcription start sites with interactions at the promoter area (striped boxes).

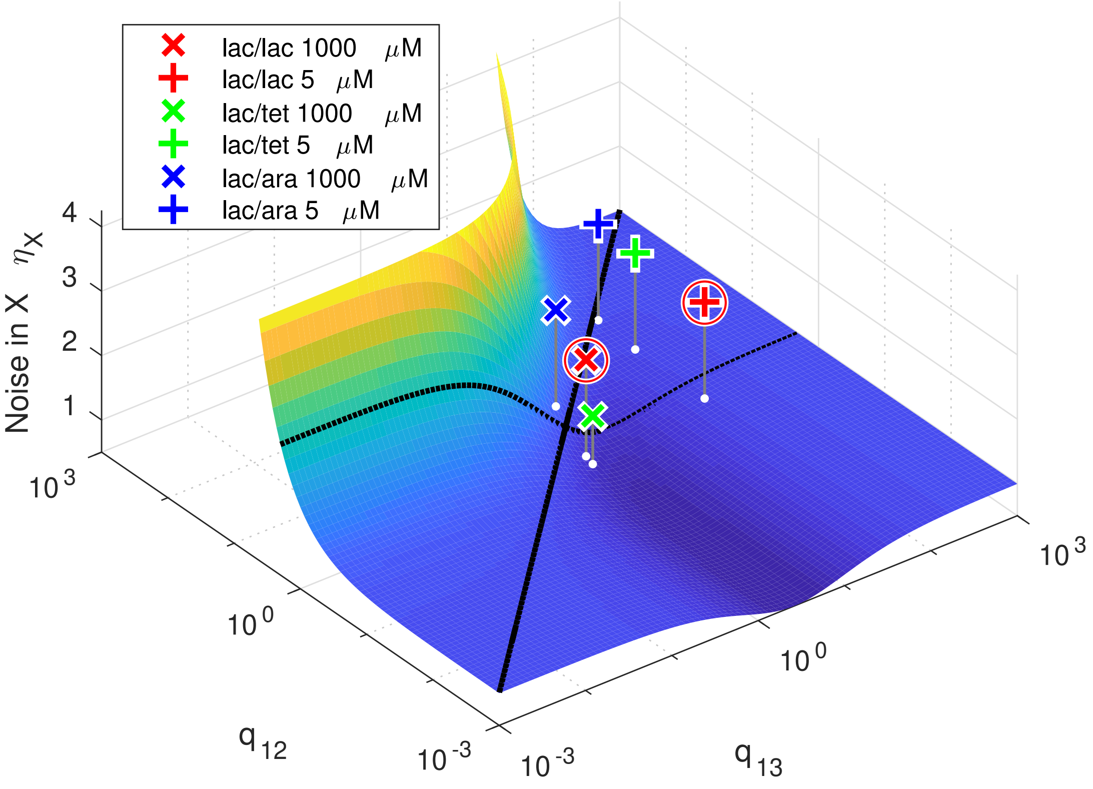


FIG. S2. Noise of RNA production of one side of a dual promoter as a function of the relative durations of closed and open complex formation and the expression ratio of the promoters. The black curves denote unity and $q_{12} = 2$ (cf. Fig. 3), and the markers the predictions for the measurements, circles representing the validated ones (cf. Table III).

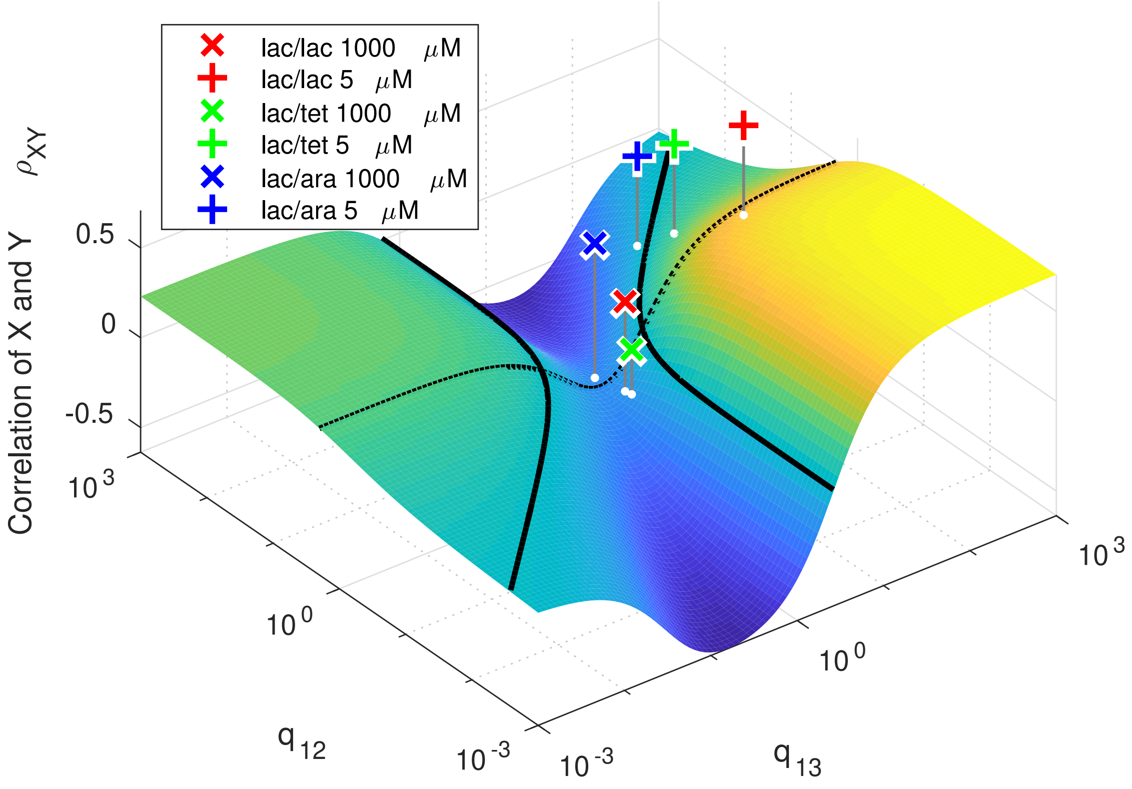


FIG. S3. Predicted correlation between the RNA productions initiated by each start site of the dual promoter as a function of the relative durations of closed and open complex formation and the expression ratio of the promoters. The black curves denote zero and $q_{12} = 2$ (cf. Fig. 4A), and the markers the predictions for the measurements.

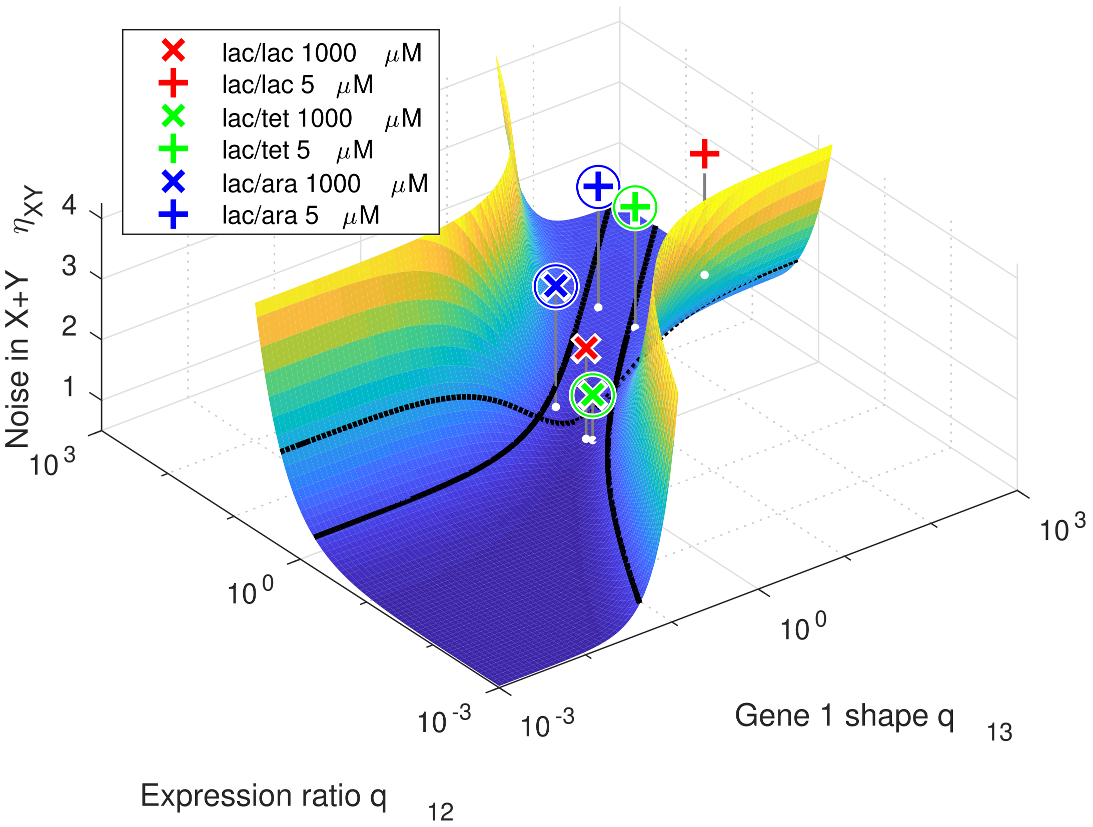


FIG. S4. Noise of RNA production of the dual promoter as a function of the relative durations of closed and open complex formation and the expression ratio of the promoters. The black curves denote unity and $q_{12} = 2$ (cf. Fig. 4B), and the markers the predictions for the measurements, circles representing the validated ones (cf. Table II and Table III).

[S1] W. R. McClure, Proc. Natl. Acad. Sci. U.S.A. **77**, 5634 (1980).

[S2] D. A. McQuarrie, J. Appl. Probab. **4**, 413 (1967).

[S3] D. T. Gillespie, Annu. Rev. Phys. Chem. **58**, 35 (2009).

[S4] R. M. Saecker, M. T. Record, Jr., and P. L. deHaseth, J. Mol. Biol. **412**, 754 (2011).

[S5] D. T. Gillespie, Physica A **188**, 404 (1992).

[S6] A.-B. Muthukrishnan, M. Kandhavelu, J. Lloyd-Price, F. Kudasov, S. Chowdhury, O. Yli-Harja, and A. S. Ribeiro, Nucl. Acids Res. **40**, 8472 (2012).

[S7] J. Lloyd-Price, S. Startceva, V. Kandavalli, J. Chandraseelan, N. Goncalves, S. M. D. Oliveira, A. Hakkinen, and A. S. Ribeiro, DNA Res. **23**, 203 (2016).

[S8] S. M. D. Oliveira, A. Hakkinen, J. Lloyd-Price, H. Tran, V. Kandavalli, and A. S. Ribeiro, PLoS Comput. Biol. **12**, e1005174 (2016).

[S9] A. Hakkinen and A. S. Ribeiro, Bioinformatics **31**, 69 (2015).

[S10] V. K. Kandavalli, H. Tran, and A. S. Ribeiro, BBA Gene Regul. Mech. **1859**, 1281 (2016).

[S11] K. M. Herbert, A. La Porta, B. J. Wong, R. A. Mooney, K. C. Neuman, R. Landick, and S. M. Block, Cell **125**, 1083 (2006).

[S12] T. Rajala, A. Hakkinen, S. Healy, O. Yli-Harja, and A. S. Ribeiro, PLoS Comput Biol **6**, e1000704.

[S13] M. H. Larson, R. A. Mooney, J. M. Peters, T. Windgassen, D. Nayak, C. A. Gross, S. M. Block, W. J. Greenleaf, R. Landick, and J. S. Weissman, Science **344**, 1042 (2014).

[S14] A. Hakkinen and A. S. Ribeiro, Bioinformatics **32**, 1346 (2016).

[S15] M. Kaern, T. C. Elston, W. J. Blake, and J. J. Collins, Nat. Rev. Genet. **6**, 451 (2005).

[S16] D. R. Cox, *Renewal theory* (Methuen, London, UK, 1962).

[S17] J. M. Pedraza and J. Paulsson, Science **319**, 339 (2008).

[S18] D. Huh and J. Paulsson, Proc. Natl. Acad. Sci. U.S.A. **108**, 15004 (2011).

[S19] M. Herbert, A. Kolb, and H. Buc, Proc. Natl. Acad. Sci. U.S.A **83**, 2807 (1986).

[S20] A. Dempster, N. Laird, and D. Rubin, J. Royal Stat. Soc. B **39**, 1 (1977).

[S21] I. Golding and E. C. Cox, Proc. Natl. Acad. Sci. U.S.A. **101**, 11310 (2004).
